# Efficient CRISPR/Cas9 genome editing in a salmonid fish cell line using a lentivirus delivery system

**DOI:** 10.1101/734442

**Authors:** Remi L. Gratacap, Tim Regan, Carola E. Dehler, Samuel A.M. Martin, Pierre Boudinot, Bertrand Collet, Ross D. Houston

**Author notes:** Correspondence: Remi L. Gratacap. Ross D. Houston.

## Abstract

Genome editing is transforming bioscience research, but its application to non-model organisms, such as farmed animal species, requires optimisation. Salmonids are the most important aquaculture species by value, and improving genetic resistance to infectious disease is a major goal. However, use of genome editing to evaluate putative disease resistance genes in cell lines, and the use of genome-wide CRISPR screens is currently limited by a lack of available tools and techniques. In the current study, an optimised protocol using lentivirus transduction for efficient integration of constructs into the genome of a Chinook salmon (*Oncorhynchus tshwaytcha*) cell line (CHSE-214) was developed. As proof-of-principle, two target genes were edited with high efficiency in an EGFP-Cas9 stable CHSE cell line; specifically, the exogenous, integrated EGFP and the endogenous RIG-I locus. Finally, the effective use of antibiotic selection to enrich the successfully edited targeted population was demonstrated. The optimised lentiviral-mediated CRISPR method reported here increases possibilities for efficient genome editing in salmonid cells, in particular for future applications of genome-wide CRISPR screens for disease resistance.

## 2 Introduction

Aquaculture is the fastest growing food production sector, and has overtaken capture fisheries as the primary source of seafood for human consumption (FAO, 2018). However, farmed production of finfish, shellfish, and crustacean species all suffer from infectious diseases that can have negative impacts on animal welfare, the environment, and on commercial viability, constraining future expansion. Selective breeding for improved disease resistance is a promising avenue to tackle these diseases, and has been widely practiced for Atlantic salmon (*Salmo salar*) and other salmonid species (Houston, 2017; Yáñez et al., 2014). Family-based selection based on siblings trait recording is now typically augmented with the use of molecular genetic markers, either via marker-assisted selection (based on markers linked to quantitative trait loci, QTL) or genomic selection using genome-wide markers to predict breeding values (Zenger et al., 2018). However, to date, little is yet known about the functional genes and variants underlying this genetic resistance to disease.

Genome editing using reprogrammed CRISPR/Cas9 systems has emerged as a revolutionary tool to make specific and targeted changes to genomes of species’ of interest. CRISPR/Cas9 can facilitate identification and characterisation of specific functional variants underlying QTL affecting the trait of interest. The technology facilitates precise gene knockout, activation or inhibition, and can also allow targeted deletion, insertion and even epigenetic modification of genomic DNA (Knott and Doudna, 2018). As such, CRISPR/Cas9 can be applied to test targeted perturbation of candidate genes and variants within QTL regions, to assess the consequence on the trait of interest. This knowledge raises the possibility of enhancing genomic selection accuracy via increased weighting on functional variants. In addition, genome editing can potentially by applied to create *de novo* variation, or to introduce favourable alleles segregating in closely related strains or species (Gratacap et al., 2019).

The aforementioned genome editing approaches typically focus on a single target locus, and there are several examples of successful CRISPR editing of single loci *in vivo* in farmed fish species (e.g. Datsomor et al., 2019; Edvardsen et al., 2014), reviewed in (Gratacap et al., 2019)). CRISPR/Cas9 has also been successfully applied in salmonid cell culture models to investigate specific components of the interferon pathway (Dehler et al., 2019). Another exciting application that has emerged in recent years is the development of genome-wide CRISPR knock out (so-called GeCKO) screens in cell culture models (Doench, 2018). This involves creation of a library of tens/hundreds of thousands of guide RNAs (gRNA) targeting every gene in the genome of the species of interest, or can also be used to target non-translated regions such as enhancers or miRNA (Fulco et al., 2019). These guides are then synthesised, packaged into a lentivirus vector, and transduced into a cell line constitutively expressing Cas9 (or alternatively the transduced construct can also code for the Cas9 protein). The lentivirus dose used results in approximately one gRNA integration per cell. The cell pool is then screened (e.g. using a pathogen challenge) and the selected cells (surviving, fluorescently labelled, or another marker of selection) sequenced. The enrichment or depletion of gRNAs compared to the control population informs on the role of their target genes in the phenotype under investigation. This approach has led to fundamental host-pathogen discoveries, particularly in virology as the cell intrinsic nature of the innate immune response is very well suited to interrogation with this platform. Such screens have led to the discovery of the Norovirus receptor (Orchard et al., 2016) and the role of endoplasmic reticulum membrane complex in Zika virus infection (Savidis et al., 2016). Genome-wide screens have been applied in several species, from humans to fly (Viswanatha et al., 2018), including parasites such as *Toxoplasma gondii* (Sidik et al., 2018), and plants *(Meng et al., 2017)* but has yet to be applied in farmed fish species, where it could have major potential for discovering genes involved in disease resistance, and improving knowledge of host-pathogen interaction.

There are currently several barriers and knowledge gaps preventing the application of GeCKO screens in fish. However, one of the major one was overcome when a potentially suitable cell line was created by Dehler et al (2016), who used a random plasmid integration event to stably express Cas9 in the Chinook salmon (*Oncorhynchus tshawytscha*) cell line (CHSE-EC), a cell line frequently used to study viral diseases of interest to farmed salmonid production. Further, this transgenic line was also modified to stably express EGFP, and Dehler et al used electroporation of gRNA to successfully knockout the integrated EGFP locus, reporting approximately 35 % successful editing. Using this approach, they developed clonal lines of CHSE STAT2 KO and explored the role of this gene in antiviral immunity (Dehler et al., 2019). Alternatively, the genome editing cargo can be delivered by transient transfection of a plasmid expressing Cas9 and gRNA. Escobar et al, (2019) successfully edited the genome in the CHSE cell line but reported a low transfection/expression efficiency (10%) and did not report editing efficiency. To harness the CHSE-EC (or similar Cas9 stable) cell lines as a platform for high-throughput screens, an efficient lentivirus delivery system for the GeCKO library would be highly desirable.

Lentiviral transduction of Cas9 and gRNA has several advantages over other methods, such as electroporation of ribonucleoprotein or transient plasmid transfection. It efficiently integrates into the genome, enabling the creation of stable cell lines, and allows for the integration of antibiotic resistance markers and fluorescent reporters to perform enrichment of edited cells. A multitude of lentivirus plasmid constructs already exist, and have been successfully tested in various species. For these reasons, lentivirus is the delivery method of choice for GeCKO libraries, but has not yet been developed in fish cell lines. In the current study, an efficient lentivirus-based method for genome editing in a salmonid fish cell line is presented. The delivery of lentivirus was optimised to allow integration of a transgene at high efficiency. Using a lentivirus delivered gRNA in the salmonid cell line CHSE-EC, a very high efficiency of genome editing was obtained, after antibiotic enrichment of cells containing both Cas9 and specific gRNA. Finally, as proof of principle, the method was applied to establish a polyclonal cell line enriched for retinoic acid-inducible gene-I-like (RIG-I) knock-out cells.

## 3 Results

### 3.1 Efficient transduction of chinook salmon cells with lentivirus

To improve CRISPR/Cas9 delivery and genome editing efficiency, lentiviral transduction was optimised for use in salmonid cell lines. This was initially performed using a fluorescent reporter construct (CMV:EGFP) in the Chinook salmon cell line (CHSE-214, referred to hereafter as CHSE). To optimise the transduction efficiency, three major variables were tested (Park et al., 2015; Phenix et al., 2000); specifically (i) the impact of incubation temperature; (ii) the impact of including a spinfection (or spinoculation); and (iii) the impact of the duration of the incubation. Flow cytometry was used to determine the efficiency of transduction by measuring the number of fluorescent cells (therefore assumed to be transduced with the CMV-EGFP), with the outputs from the optimised settings shown in Figure 1a and 1b.

**Figure 1.**
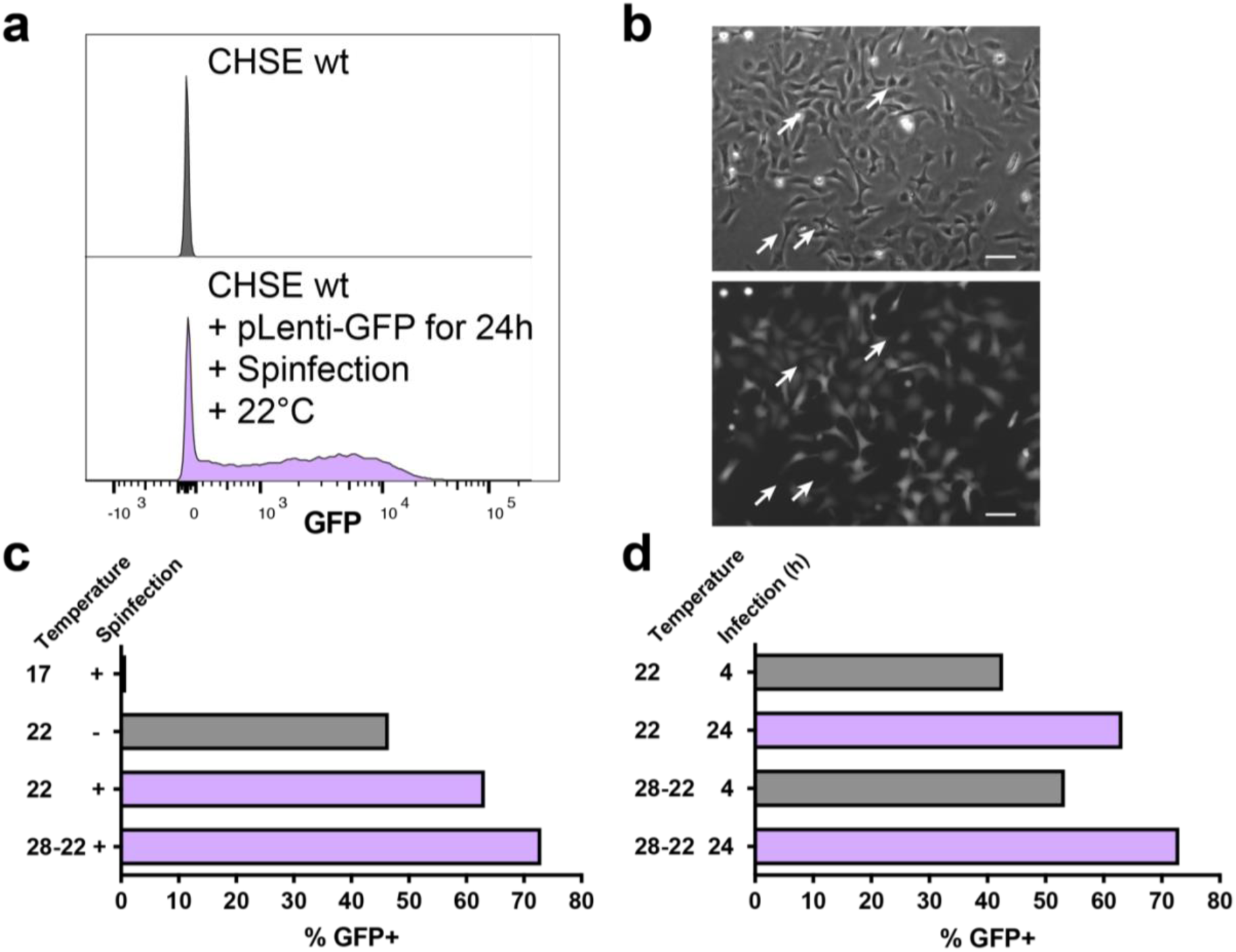
CHSE salmon cells are efficiently transduced with lentivirus. **a**, Efficient transduction of CMV:EGFP in CHSE Chinook salmon cell line by lentivirus. Salmon cells were spinfected at 22 °C with neat lentivirus supernatant for 2h at 1,000 × g. The cells were incubated for 24 h and the media was replaced. After 2 weeks of expansions, fluorescence was recorded by flow cytometry using CHSE wt (not transduced) as control. Split histogram of control cells (top) and pLenti-GFP transduced CHSE with optimal conditions (bottom). Data were normalised to Mode (relative percentage of cells rather than number). **b**, Representative image of CHSE cells 8 days post transduction with pLenti-GFP. Differential Interference Contrast (top) and GFP (bottom) channels are represented. Arrows point to GFP negative cells. Scale bar 20 µm. **c-d**, Different conditions of the optimisation protocol with optimal conditions in purple. **c**, Incubation temperature and spinfection influences the efficiency of transduction. Cells were transduced with neat lentivirus supernatant, spinfected for 2 h at 1,000 × g (or not) and incubated at 17 °C (17), 22 °C (22) or heat shocked for 4 h at 28 °C followed by 22 °C for 2 weeks (28-22) until flow cytometry. **d**, Incubation of the cells with the virus for 4h reduces the efficiency of transduction. CHSE were transduced with neat lentivirus dose at different temperatures and media was changed after 4 or 24h.

Temperature of incubation had a major impact on transduction efficiency, with an increased transduction efficiency from 1 % to 63 % as the temperature was increased from 17 °C to 22 °C (Fig1c). Further increase in temperature to 28 °C for 4 hours (heat shock) followed by 22 °C incubation (denoted 28-22) resulted in a minor increase in transduction efficiency, from 63 to 73 % (Fig1c), but overnight incubation at 28 °C caused mortality in the CHSE cells (data not shown). Using the settings of 22 °C incubation for 24 hours and neat supernatant of lentivirus, we included a spinfection step and show an improvement of transduction efficiency from 47 % to 63 % (Fig 1c). Finally, reducing the incubation time of the cells with the lentivirus from 24 h to 4 h reduced the transduction efficiency from 63 % to 43 % for 22 °C incubation and 73 % to 53 % for 28-22 incubation group (Fig 1d). Therefore, we propose an optimised protocol for efficient transduction of CHSE cells using neat lentivirus supernatant on suspended cells, together with a spinfection step (2h at 1000 × g) and incubation for 24 h at 22 °C (Figure 1a, 1b). These optimised settings were used for downstream experiments.

### 3.2 Efficient editing of the Chinook salmon genome using a lentivirus system

After optimising the transduction conditions by integrating a GFP expressing construct into the CHSE cell line, a modified CHSE cell line stably expressing Cas9 and EGFP (CHSE-EC (Dehler et al., 2016)) was used to test Cas9-mediated genome editing in these cells. A plasmid, containing a human U6 promoter to drive the expression of a second-generation gRNA scaffold and the puromycin antibiotic resistance gene was used (Tzelepis et al., 2016). The use of the human U6 promoter to drive the expression of a gRNA has previously been reported to be inefficient in this salmonid cell line (Escobar-Aguirre et al., 2019). Therefore, to first test this, the gRNA sequence for EGFP (Sanjana et al., 2014) was cloned in this plasmid (pKLV2_EGFP), and transduced CHSE-EC cells using the optimised protocol described previously.

Neat lentivirus supernatant was used to transduce CHSE-EC, which resulted in up to 89.9 % reduction in fluorescent cells by flow cytometry (Fig2a, b). A 10-fold dilution of the lentivirus supernatant was also used to validate the antibiotics enrichment. This yielded a 42.9 % loss of fluorescence, which could be effectively enriched using puromycin selection for 1 week, to 82.5 % loss (Lo + Puro, Fig2b, c). In close agreement with the flow cytometry data, Sanger sequencing analysis of the target region showed 70 % editing (cutting efficiency using ICE, Synthego Inc) of the cells in the neat supernatant group (and 58 % in the low dose enriched, Fig2c). Deconvolution of the chromatogram edit patterns (ICE analysis) shows that the majority of the edited sequences had a 1 bp insertion (27 %) and only 15 % of the sequences were estimated to be unedited (Fig 2d).

**Figure 2.**
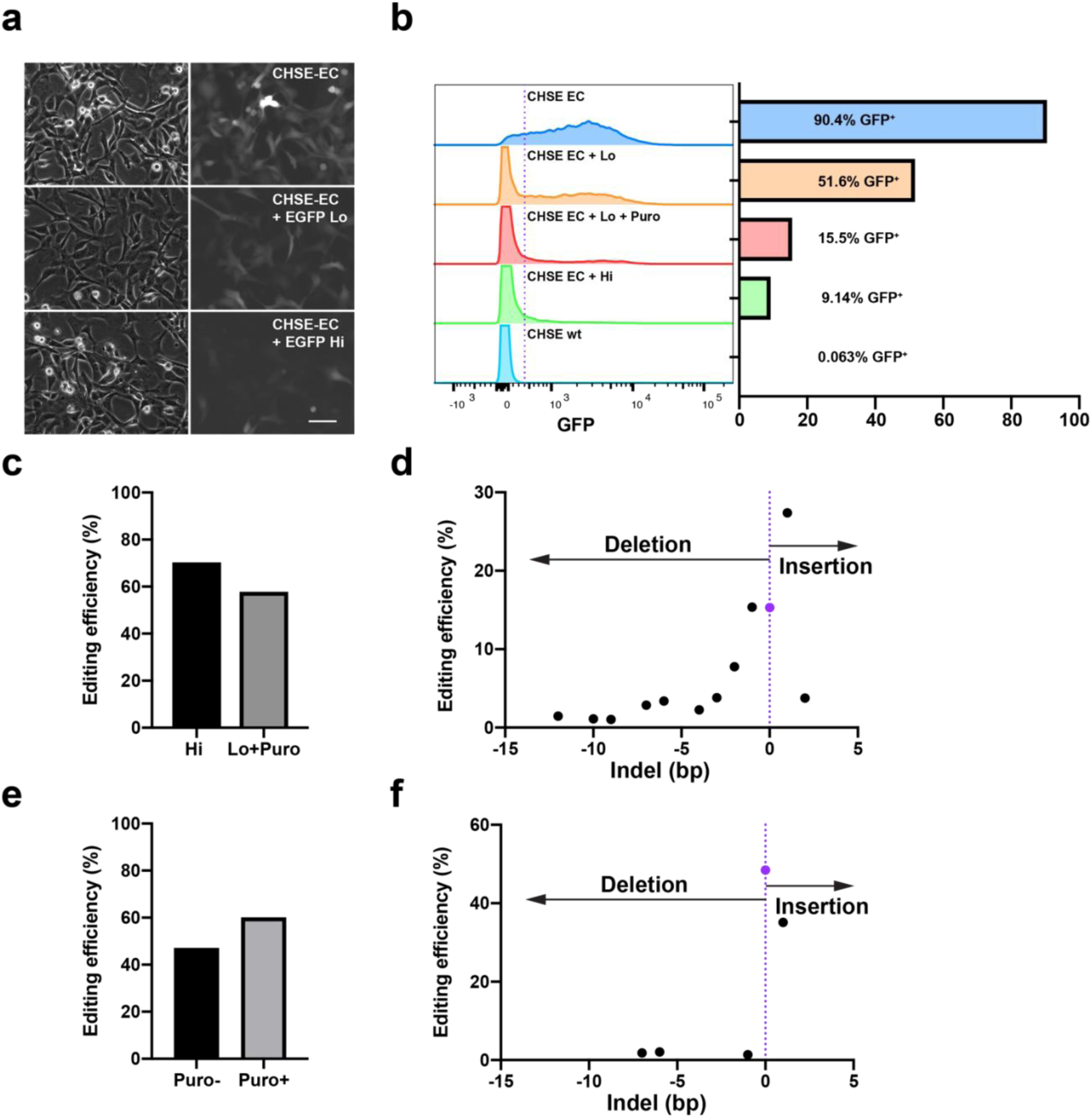
The genome of CHSE-EC salmon cells is efficiently edited with lentivirus. **a-d**, Efficient editing of EGFP in CHSE-EC Chinook salmon cell line by lentivirus. CHSE-EC cells were spinfected for 2 h at 1,000g with neat (Hi) and 1:10 dilution (Lo) of lentivirus supernatant and incubated at 22 °C. After 2 weeks of expansion, puromycin was added to the Lo group (0.25 ug / mL, for 1 week, Lo + Puro). Fluorescence was imaged by epifluorescence microscopy (**a**) and recorded flow cytometry (**b**) using CHSE wt and CHSE-EC (not transduced) as controls. Split histogram of fluorescence and corresponding proportion histogram of control cells (top and bottom) and CHES-EC transduced with high and low concentration (with and without puromycin treatment) lentivirus supernatant. Scale bar in a represents 20 µm. **c**, Genome editing efficiency in CHSE-EC by Sanger sequencing of PCR amplified product of the target loci, estimated by ICE analysis. **d**, Detail of indels frequency in EGFP edited Hi group (from panel c), estimated by ICE analysis. Purple dot denotes the unedited sequence (0 bp). **e-f**, Efficient editing of RIG-I in CHSE-EC. **e**, Genome editing of the RIG-I gene in CHSE-EC. CHSE-EC cells were transduced similar to EGFP targeting (Puro-) and selected with puromycin (Puro+) and efficiency estimated by ICE analysis of Sanger sequencing chromatogram from the PCR amplified target region. **f**, Detail of indels frequency in RIG-I edited Puro-group (from panel e), estimated by ICE analysis. Purple dot denotes the unedited sequence (0 bp).

In summary, the results show that the pKLV2 plasmid (containing the human U6 promoter) is functional and effective at transcribing gRNA in the CHSE cell line, and that the lentivirus delivery strategy (either neat, or diluted together with antibiotic selection for enrichment) leads to very high genome editing efficiency.

To validate that the editing strategy was efficient on a salmon gene, the Chinook salmon retinoic acid-inducible gene-I-like (RIG-I) locus (DDX58, NCBI Gene LOC112222314) was targeted, by creating a pKLV2_RIG-I construct. This gene was chosen as it is central in the antiviral defense against RNA virus (Yoneyama et al, 2004), it exists as a single copy in the chinook salmon genome, and because knockout cell lines may be useful models for future studies of host response to these viruses. The CHSE-EC cells were transduced with pKLV2_RIG-I supernatant, followed by puromycin selection for 7 days. At least 47% of the cells were edited at the desired locus (Puro-, Fig2e) and this was enriched to 60 % using puromycin selection (Puro+, Fig2e). The pattern of indels resulting from the CRISPR editing was similar to EGFP edited cells, with 1 bp insertions being the most common mutation arising from the editing (Fig2f). This demonstrates that endogenous genes in the chinook salmon CHSE cell line can be targeted for genome editing using lentivirus.

Taken together, these results show that the lentivirus delivery strategy described herein, together with the Cas9 expressing CHSE cell line (CHSE-EC) can easily, rapidly and efficiently be used to edit the genome of salmonid cells, and suggests that the method could be applied to the new generation of CRISPR/Cas9 platforms such as base editors and CRISPR activation/inhibition.

## 4 Discussion

The CHSE-214 salmonid cell line can be efficiently transduced with lentivirus; the first demonstration of lentivirus-mediated editing in salmonid fish cells. The optimised protocol described results in the efficient genome editing of a salmonid cell line to date. This study paves the way to high-throughput genome editing in salmonid cells, and for simpler generation of transgenic cell lines.

Retroviruses have widely been used to infect animal cells and to deliver cargo which can be integrated or not in the genome (Cronin et al., 2005). Over 25 years ago, it was shown that fish cells (zebrafish, *Danio rerio* ZF4, Rainbow trout, *Oncorhynchus mykiss* RTG-2 and Chinook salmon, CHSE-214) could be transduced with retrovirus (Moloney murine leukemia virus, MLV) after subtyping it with the VSV-G envelope glycoprotein (from Vesicular Stomatitis Virus) (Burns et al., 1993). Thereafter, several viral delivery vectors have been tested for the expression of genes (integrative or non-integrative system) in several fish cells and *in vivo*, including MLV (Liu et al., 2015), Semliki Forest Virus (SFV) (Phenix et al., 2000), lentivirus, adeno-virus (Ad) and adeno-associated virus (AAV) (Kawasaki et al., 2009; Overturf et al., 2003; Sarmasik et al., 2001), with levels of efficiency varying from very high [Ad in zebrafish cells (Kawasaki et al, 2009)] to low [MLV in medaka stem cells (Liu et al, 2015)]. Therefore, the transduction of cell lines has been shown to be efficient with the retrovirus system, but its adoption to edit the genome of fish cell lines has not been demonstrated.

In the present study, the delivery method of the retrovirus-derived second-generation lentivirus system (Naldini et al., 1996) was optimised to stably integrate an EGFP construct in the salmonid cell line CHSE-214. Transduction efficiency is dependent on temperature and is almost completely blocked at incubation below 17 °C. The cell type and temperature influence on lentivirus transduction efficiency has been reported previously in mammalian cells (O’Neill et al., 2010; Park et al., 2015). In fish cell lines, SFV infection was reduced 1000-fold in the Atlantic salmon F95/9 cell line and completely blocked in CHSE-214 as the temperature was decreased from 25 to 15 °C (Phenix et al., 2000). This temperature restriction is fortunately not a limiting issue as most fish cell lines, derived from cold or warm water species, are adapted to temperatures of 20 °C or above (Ott, 2004). Similar to several studies in mammalian cells, adding a spinfection step (1000 × g for 1-2 h) greatly enhanced the transduction efficiency of LV in the current study (O’Doherty et al., 2000; Park et al., 2015). The relative ease of stably integrating a reporter construct and possibly tagging the native protein (Serebrenik et al., 2019) is likely to help with *in vitro* research in salmonid cell lines.

A major advantage of lentivirus transduction is the possibility to integrate an antibiotic resistance cassette to enrich the transduced cells. Using this approach, it is possible to use a relatively low viral dose, which ensures that a single copy of the insert is integrated in the genome, and subsequently enrich the transduced cells with antibiotic treatment to very high proportion of cells. The high transduction efficiency and genome editing achieved offer advantages over less efficient gene editing methods such as the possibility to directly use the cells generated as a pooled population without the tenuous process of single cell cloning. However, it should be noted that lentivirus, like most integrating retroviruses, tend to integrate at the site of active transcription (active genes) and that an immune response (interferon stimulated genes) has been reported with lentivirus transduction (i.e. infection) (Pichlmair et al., 2007; Schröder et al., 2002).

By using the CHSE-EC cell line which constitutively expressed Cas9, in combination with a lentivirus packaging a gRNA construct targeting EGFP or RIG-I, the genome of the salmonid cells was edited at very high efficiency. Using this strategy, the previously reported editing efficiency of 35 % by electroporation of the gRNA (Dehler et al., 2016) was improved to almost 90 % editing of EGFP (as assessed by flow cytometry). Nonetheless, it should be noted that electroporation of difficult-to-transduce cells has been extremely successful and is also one of the main approaches to edit the genome of mammalian immune cells (Kim et al., 2014).

The expected positive correlation was observed between the editing efficiency estimated by Sanger sequencing and the phenotype observed when targeting the exogenous EGFP gene in the CHSE-EC salmonid cell line. However, disrupting specific pathways by precise targeting of endogenous genes might be more challenging, due to the recent salmonid whole genome duplication (approx. 95 million years ago, (Macqueen et al., 2017)). As such, it will be necessary in many cases to target all paralogs to achieve the desired phenotype. Moreover, several mechanisms have been hypothesized to explain genetic robustness (i.e. a mutation not exhibiting the expected phenotype), such as functional redundancy (Tautz, 1992), rewiring of genetic networks (Barabási and Oltvai, 2004), adaptive mutations (Teng et al., 2013) or transcriptional adaptation (El-Brolosy et al., 2019).

The optimised protocol described in this study allows for relatively fast and cost-effective gene editing in Chinook salmon cells to generate reporter or CRISPR KO lines. Cell lines from other salmonid species such as Atlantic salmon might be readily transduced with similar lentivirus systems. However, salmonid cell lines are generally slow growing and difficult to transfect with plasmids, with the CHSE being the most amenable to clonal expansion. Therefore, development of embryonic cell lines from other salmonid species may be a useful approach to generate appropriate cell lines for CRISPR KO experiments in more commercially relevant aquaculture species.

The prospect of developing and applying a genome wide CRISPR KO screen in fish cell lines is very attractive for many reasons (Doench, 2018; Gratacap et al. 2019) and the present system paves the way towards such a platform for salmonids. This has the potential to be transformative for the testing of candidate disease resistance genes generated by Genome-Wide Association Studies as well as *de novo* discovery of genes involved in host-pathogen interactions in fish.

## 5 Materials and Methods

### 5.1 Cell culture

CHSE cell line from Chinook salmon was obtained from ECACC (91041114). CHSE-EC were previously described (Dehler et al., 2016). Both salmonid cell lines were cultured in L15 media (Sigma L1518) supplemented with 10% heat inactivated fetal bovine serum (FBS, Gibco) and 1x Pen/Strep (P/S) antibiotics (Gibco). Cells were cultured at 22 °C (or see optimisation for other temperatures) without CO_2_ and split 1:4 at 80% confluency. x293T (Clontech) were cultured in DMEM + 10% FBS, Penicillin, Streptomycin and Glutamine, Sodium Pyruvate and NEAA (all from Gibco). Cells were cultured at 37 °C with 5% CO_2_ and split 1:20 at 80% confluency. Puromycin (Cayman chemicals) was used for antibiotic selection at 0.25 µg/mL for 7 days. The media was changed every 3-4 days.

### 5.2 gRNA design

All gRNAs were designed using CRISPOR (www.crispor.tefor.net) and the Chinook salmon reference genome (NCBI GCF_002872995.1, Otsh_v1.0) for off-target evaluation. The oligos and PCR primers spanning the edited locus (400-600 bp), synthesised by IDT or Sigma can be found in Table1. All oligos (Table 1) were resuspended to 100 µM in 1x TE.

**Table 1:**
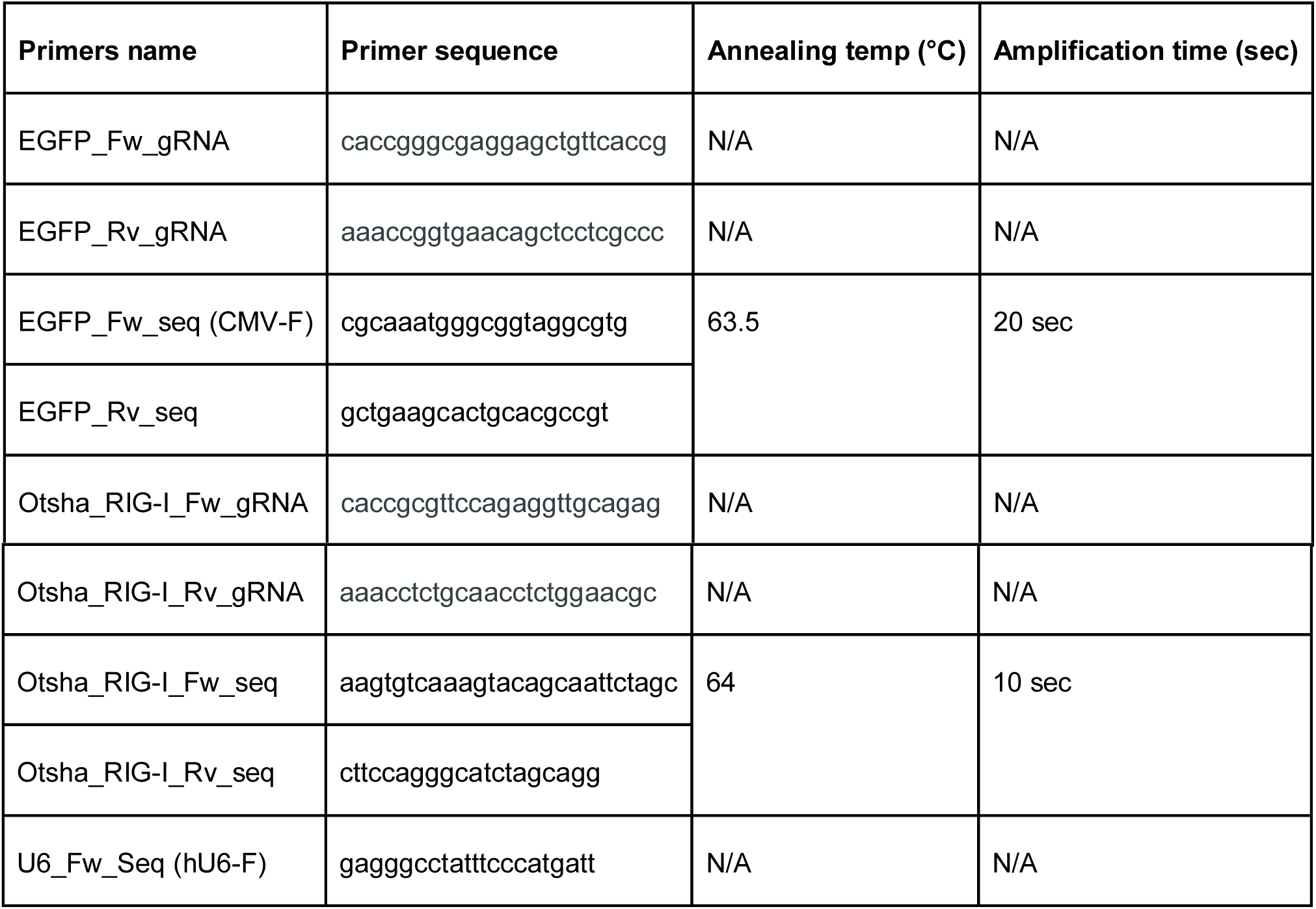
Primers used in the study (from IDT or Sigma). gRNA target sequence underlined.

### 5.3 Plasmids constructs (Table 2)

pKLV2-U6gRNA5(BbsI)-PGKpuro2ABFP-W (pKLV2) was a gift from Kosuke Yusa (Addgene plasmid # 67974). Specific gRNA for EGFP and RIG-I (Table2, underlined) were cloned according to Golden Gate reaction from ZhangLab protocol (Addgene SAM library sgRNA cloning protocol) using BbsI-HF (NEB) and Stbl3 competent *E. coli* cells (Thermo Fisher) for transformation. Briefly, 1 µL of each oligo (100 µM) were mixed with 1 µL of T4 ligation buffer (NEB) and 7 µL of water. The mix was heated to 95 °C for 5 min and cooled to room temperature. 1 µL of the annealed oligos was mixed with 25 ng of pKLV2, 1 µL of T4 ligation buffer, 9 µL of water and 0.5 µL of BbsI enzyme and 0.5 µL of T4 ligase enzyme (200 U, NEB). The mix was incubated for 10 cycles of 5 min at 37 °C and 5 min at 23 °C. Five µL were transformed in 50 µL competent *E. coli* (Stbl3, Thermo fisher). Four colonies were picked and grown in LB + Carbenicillin, the plasmids purified (NEB Monarch Plasmid miniprep) and sequenced to verify correct insertion using U6_Fw_seq primer (Table 1).

**Table 2:** Plasmids used in this study (from Addgene, USA).

| Plasmid construct | Plasmid name (Addgene) | Plasmid number (Addgene) |
| --- | --- | --- |
| pLenti CMV:EGFP_PGK:Puro | pLenti CMV GFP Puro (658-5) | 17448, (Campeau et al., 2009) |
| pLenti hU6:gRNA_PKG:Puro | pKLV2-U6gRNA5(BbsI)-PGKpuro2ABFP- | 67974, (Tzelepis et al., 2016) |

### 5.4 Lentivirus production

Supernatants containing lentiviral particles were produced by transient transfection of x293T cells using Lipofectamine 2000 (Invitrogen). To transfect one well of a 6-well plate, 1.5×106 x293T cells were plated in 4 mL of complete media and incubated overnight at 37 °C (70% confluency at 24h). The following day, 1.8 µg of lentiviral vector, 1.8 µg of psPAX2 (gift from Didier Trono, Addgene plasmid # 12260) and 0.4 µg of pMD2.G (gift from Didier Trono, Addgene #12259) were added to 160 µL of Opti-MEM (no phenol red, Gibco), 12.5 µL of Lipofectamine 2000 was added to 140 µL of Opti-MEM and both tubes incubated for 5 min at room temperature. The Lipofectamine 2000 mixture was then added to the plasmids mixture and further incubated for 20 min at room temperature. The transfection complex (300 µL) was added dropwise to one well of x293T and incubated for 48 h at 37°C with CO_2_. Viral supernatant was then harvested, centrifuged at 500 × g for 4 min, filtered using a 0.45 µm low protein retention syringe filter (Sartorius) and 0.5 mL aliquots were stored at −80 °C.

### 5.5 Lentiviral transduction

For lentiviral transduction of CHSE or CHSE-EC, 50 µL of cells (4.105 cells/mL) were mixed with 100 µL of the lentiviral supernatant (at various dilutions) in a 96-well plate or 200 µL of cells plus 400 µL of viral supernatant in 24-well plate, and centrifuged for 2h at 1,000 × g. Media was changed after 4h or 24h. After transduction, cells were incubated at the indicated temperatures and split as needed.

### 5.6 Flow cytometry

Transduced cells were detached with trypsin, centrifuged and resuspended in PBS. The cells were kept on ice and flow cytometry was performed using a Fortessa-X20 (BD Biosciences). Events were gated to remove doublet cells and the GFP intensity of each cell was analysed with FlowJo 10.6.0 (Becton Dickinson).

### 5.7 Sequencing

Genomic DNA was extracted with QuickExtract buffer (Lucigen) by adding 30 µL to a single well of a 96-well plate and incubating for 5 min. The samples were then processed according to the manufacturer’s instructions (65 °C for 6 min and 98 °C for 2 min). PCR was performed with 50 µL reactions using NEB Q5 and 1 µL of the gDNA for 33 cycles amplification at optimal annealing temperature (see Table1). Amplified products were purified using AMPure XP magnetic beads (Agencourt) according to the manufacturer’s instructions. Purified PCR products were sequenced by GATC/Eurofins. Sequence (ab1 files) were analysed with Interference of CRISPR Edits software (www.ice.synthego.com, (Hsiau et al., 2018)) to determine the editing and knock-out (KO, frame-shift or early stop codon) efficiency, compared to controls (non-edited).

## 9 Author Contributions

RH, TR and RG conceived and designed the study; RG performed the experiments; CD, SM, BC and PB contributed data and reagents; RH, TR and RG wrote the paper. All authors read and contributed to the final manuscript.

## 11 Acknowledgments

We would like to acknowledge colleagues within Ross Houston’s laboratory and Spring Tan for fruitful discussions as well as the Roslin Institute Bioimaging and Flow cytometry consortium.

